# Protein-protein interaction prediction for targeted protein degradation

**DOI:** 10.1101/2022.02.25.481776

**Authors:** O. Orasch, N. Weber, M. Müller, A. Amanzadi, C. Gasbarri, C. Trummer

**Affiliations:** Celeris Therapeutics GmbH, Salzamtsgasse 7, 8010 Graz, Austria

**Keywords:** protein-protein interactions, targeted protein degradation, ternary complex, deep graph representation learning

## Abstract

Protein-protein interactions (PPIs) play a fundamental role in various biological functions; thus, detecting PPI sites is essential for understanding diseases and developing new drugs. PPI prediction is of particular relevance for the development of drugs employing targeted protein degradation, as their efficacy relies on the formation of a stable ternary complex involving two proteins. However, experimental methods to detect PPI sites are both costly and time-intensive. In recent years, computer-aided approaches have been developed as screening tools, but these tools are primarily based on sequence information and are therefore limited in their ability to address spatial requirements and have thus far not been applied to targeted protein degradation.

Here, we present a new deep learning architecture based on the concept of graph representation learning that can predict interaction sites and interactions of proteins based on their surface representations. We demonstrate that our model reaches state-of-the-art performance using AUROC scores on the established MaSIF dataset. We furthermore introduce a new dataset with more diverse protein interactions and show that our model generalizes well to this new data. These generalization capabilities allow our model to predict the PPIs relevant for targeted protein degradation, which we show by demonstrating the high accuracy of our model for PPI prediction on the available ternary complex data. Our results suggest that PPI prediction models can be a valuable tool for screening protein pairs while developing new drugs for targeted protein degradation.

## 1 Introduction

Tackling diseases is one of humankind’s most long-lasting challenges. An essential part of this effort concerns targeting pathogenic proteins (e.g., misfolded proteins in CNS diseases or mutated proteins in cancer). Since Fischer’s “Lock and Key” model [1], drug discovery has followed the occupancy-driven drug discovery paradigm by focusing on designing molecules capable of fitting the active site of the target protein. This binding leads to inhibition and subsequent loss of the protein of interest (POI) function. Although this approach has been successful in many cases, it is limited by the fact that less than 20% of the human proteome is inhibitable, i.e., most POIs do not contain binding sites that compounds can occupy [2, 3, 4]. To address this shortcoming and expand the druggable space, new therapeutic modalities such as gene knockout (CRISPR-Cas9) [5] and gene expression knockdown (RNAi) [6] have been developed. An alternative lies in proximity-inducing compounds (PICs), which retain the advantages of small molecules (e.g., oral bioavailability and ease of production, [7]). PICs allow inducing proximity between POIs and proteins driving various cellular processes [8] such as phosphorylation [9], dephosphorylation [10], deacetylation [11] and protein degradation [12], allowing to repurpose the elaborate machinery available in every cell.

One mode of PIC-based drug design is targeted protein degradation (TPD) with bifunctional degraders, first described in the early 2000s [12]. The degraders used for TPD are small molecules that induce proximity between a POI and E3 ligase complex (Figure 1A). This leads to the passage of ubiquitin onto the POI, which triggers its subsequent degradation by the ubiquitin-proteasomal system (Figure 1B; see [13]). TPD, therefore, results in the selective degradation of proteins, accomplished by hijacking existing degradation pathways in human cells. This approach has several advantages [14, 15] such as allowing to target proteins that are not inhibitable by conventional means, and thus has become a significant research focus within the field of drug discovery [16, 15, 17].

**Figure 1:**
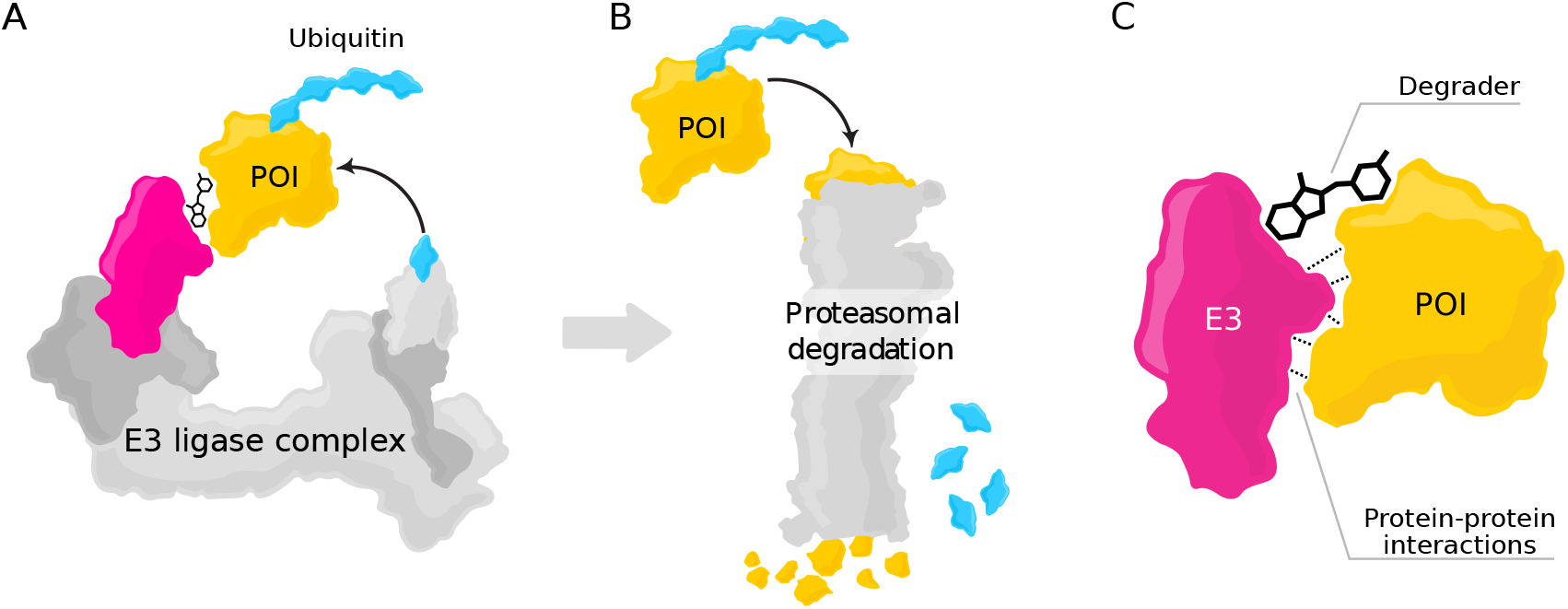
The relevance of protein-protein interactions for targeted protein degradation with bifunctional degraders. (A) To accomplish targeted protein degradation, protein of interest (POI, yellow) is linked to the receptor (magenta) of an E3 ligase complex via a degrader molecule. Together, POI, degrader, and E3 ligase form a ternary complex, allowing the passage of ubiquitin to the POI. (B) Ubiquitination of the POI leads to its degradation via the proteasomal system. The ubiquitin is recycled. (C) While the degrader molecule is instrumental for bringing the two proteins into proximity, cooperativity between the E3 ligase and the POI – i.e., strong protein-protein interactions – is essential for the formation of a stable ternary complex.

The development of new drugs operating with TPD requires predicting the interactions between the E3 ligase and the POI. Strong protein-protein interactions (PPIs) are required for the formation of a ternary complex (E3 ligase – degrader – POI, see Figure 1C), a prerequisite for POI ubiquitination [18]. If the two proteins do not interact favorably, the degrader molecule will not induce the formation of a ternary complex. In this case, for a particular POI, a different E3 ligase with favorable interactions might be considered instead (commonly used E3 ligases include, among others, CRBN and VHL [19]) if reliable PPI prediction methods are available.

There are different approaches to studying and predicting PPIs [20]. Experimental techniques are reliable for detecting structures of small proteins or single monomers, but they have intrinsic limits regarding protein complexes (see, e.g., [21, 22]) and are time- and resource-intensive [23]. To overcome these limitations, computational methods to predict 3D protein complexes and the corresponding PPIs have been developed. Many current methods rely on machine learning and can be classified into one of two groups based on the available input: template-free and template-based approaches [23]. Template-free methods provide predictions based on the peptide sequence only. While current machine learning-based template methods for protein structure and PPI prediction have shown promising results [24], they are often computationally costly, limiting their use when many inputs need to be evaluated [23]. On the other hand, template-based (e.g., [25]) approaches compute their outputs using 3D structural information as input. While these methods currently provide the most accurate predictions, a limitation lies in the need for the 3D data (for training and prediction), which is not always available [26]. The scarcity of data is particularly grave for ternary complexes in the context of TPD, where at the time of this writing, only sixteen structures are available in the Protein Data Bank [27].

This work presents a new framework for PPI prediction and shows its usefulness for predicting the interactions within the available ternary complex data. Our model processes proteins via their surface mesh [28] using deep graph representation learning (DGRL, see [29]), a popular approach of processing graph-based information such as molecular structures, which has been shown to result in superior models within the field of computational drug discovery [30]. We overcome the problem of limited training data by employing transfer learning [31]: for training our model, we use data from ordinary PPIs (available at a larger scale) and use the trained model to predict the PPIs within ternary complexes. We show that our model is competitive with the current state-of-the-art template-based PPI prediction method [25]. We furthermore introduce a new PPI dataset including more diverse data than the commonly used MaSIF dataset [28], and show that it is essential for capturing the broad range of PPIs present in ternary complexes. Our results suggest that PPI prediction can predict the PPIs underlying new ternary complexes and thus may serve as an essential tool in the development of new drugs for TPD.

## 2 Methods

This section describes the details of our model architectures, which map 3D atomic protein structures from the Protein Data Bank (PDB [27]) to binary outputs. We consider two PPI prediction tasks (see [25]):

- *Binding site prediction*: one PDB file is used as model input, and the binary output describes whether a particular location on the protein surface constitutes a possible site for protein interactions (see Figure 2A).
- *Interaction prediction*: two PDB files are processed, and the binary output describes whether the two proteins interact at a particular site (see Figure 2B).

**Figure 2:**
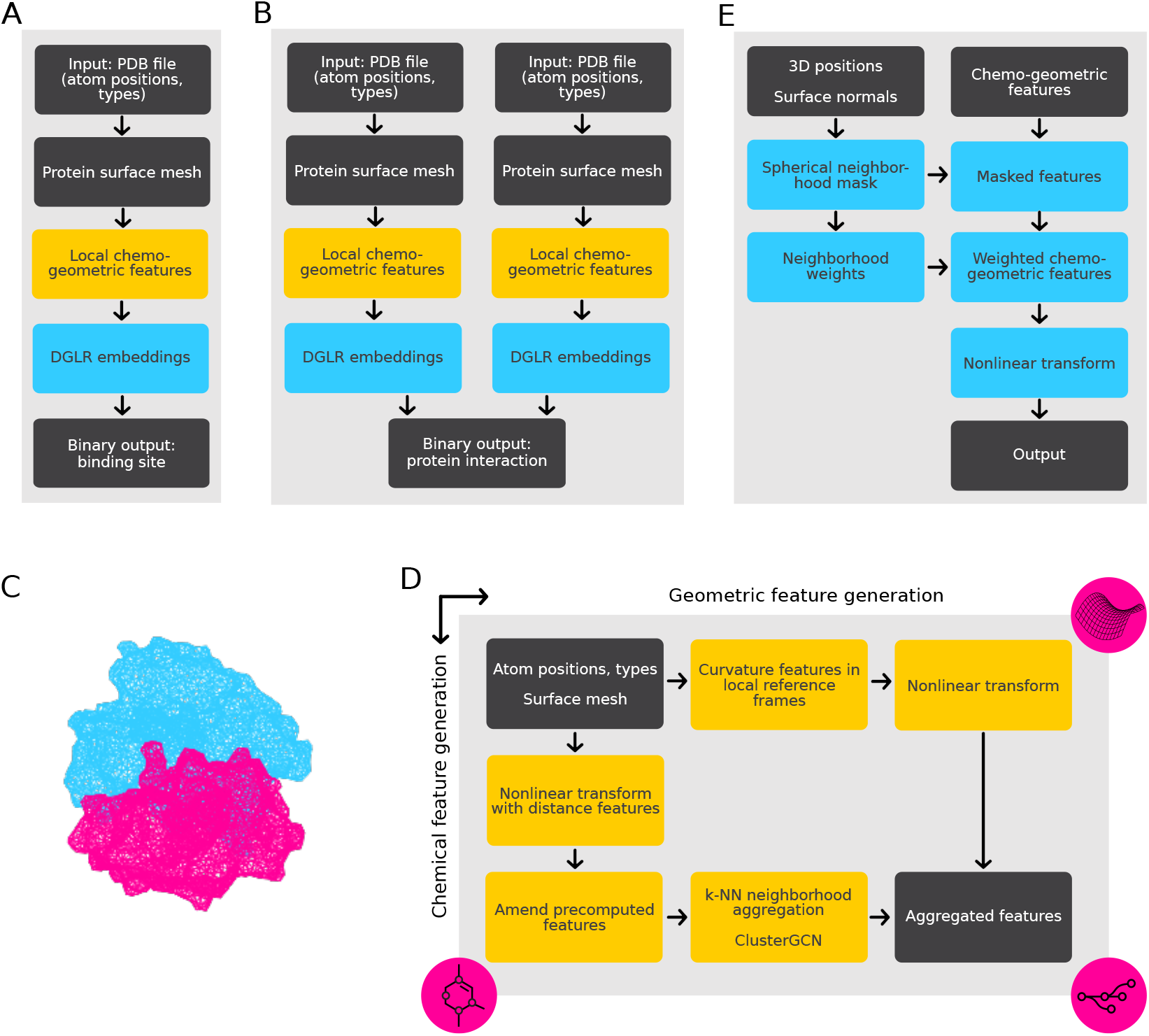
Model details. (A) Overall workflow for binding site prediction. The model outputs binary predictions of site activity for different points on the surface of proteins which are input via PDB files. The main processing steps are surface mesh generation followed by the computation of local chemo-geometric features, which are processed in a DGRL pipeline. (B) Overall workflow for interaction prediction of two proteins. Each protein is processed separatedly in a pipeline similar to that used for binding site prediction. The final processing step combines the learned features and produces a binary output. (C) Example of meshed protein surfaces (PDB entry 1A01, computed with EDTSurf [32, 33]). (D) Details of chemo-geometric feature generation. Chemical and geometrical features are generated in separate streams. The geometrical feature generation (horizontal) consists of a learned embedding of curvature features estimated in the neighborhood of a point under consideration (see text for details). The chemical feature computation (vertical) includes learned embeddings with added distance-dependend and precomputed chemical features, which are aggregated within a neighborhood and processed with a single elementary Cluster-GCN layer [34, 35] (see text for details). (E) Details of final DGRL processing. The chemo-geometric within a spherical neighborhood around a point under consideration influence the features at this point using learned weights, generated from 3D positions and surface normals. The weighted features are processed with a multilayer perceptron (MLP), resulting in the final output.

In both cases outputs are generated for various locations on the protein surface. Since our models for these two tasks share most of their components, we first provide a high-level overview before describing the components in detail.

### 2.1 Model overview

The overall model architectures for both prediction of interaction sites (Figure 2A) and protein interaction prediction (Figure 2B) are very similar, as in both cases, the input consists of proteins and the output consist of a binary value. In particular, the initial processing sequence of protein surfaces estimation, chemo-geometric feature generation, processing with DGRL are identical. The differences lie in the final processing layers, where we learn the binary classification specific to one of the two tasks: interaction site prediction (requiring input from only a single protein) or the prediction of PPIs (requiring two inputs). Additionally, the weights of the trainable components within the two pipelines are not identical and are trained separately for their respective tasks. We furthermore performed separate hyper-parameter tuning for the two different cases.

Our model uses a deep graph-representation learning (DGRL) pipeline to process the structural information of the proteins using graph representations. The input is based on surface meshes (Figure 2C), which are commonly considered to be suitable graph representations for proteins [28]. We use this representation together with domain-specific features (Figure 2D) and process them using graph-based convolutional neural networks (Figure 2E), which are engineered to respect the geometric and chemical properties of the biomolecules. In the graph-based representation, nodes within a neighborhood share common properties, thus allowing the model to reproduce real-world effects of local molecular interactions. Such relations, which are expressed with edges in the graph, can be “summarized” with the help of weight sharing. This allows our pipeline to learn expressive features that accurately discriminate protein-protein interactions.

### 2.2 (Pre-)Processing of 3D Structures into graph representations

The first processing step maps the initial 3D protein structures from PDB to a suitable surface representation. We use meshes created by EDTSurf [32, 33], where the *minimal macromolecular surface* of the underlying atomic point cloud is computed using a Euclidean distance field. Hence, the relevant graph data structures are the virtual surfaces represented by the mesh vertices. The edges are generated on the fly when required (see below).

### 2.3 Chemo-geometric feature generation

Our model learns expressive chemo-geometric features based on 3D atom coordinates, atom types, surface representation, and selected precomputed features (e.g., hydrophobicity) of the protein at each surface location under consideration. These features are fed to different neural network architecture, optimized to learn the most suitable chemical and geometric embeddings (Figure 2D). The results are concatenated and used as input to the DGRL model. In the following, we describe the details of chemical and geometric feature generation.

#### 2.3.1 Chemical features

Since we are working with surface representations of proteins, it is necessary to aggregate the near-surface atomic information into abstract feature vectors assigned to the mesh vertices. We, therefore, embed the chemical features using a k-NN directed graph where the surface point is connected to k of its underlying atoms. The chemical information associated with the atoms is processed via DGRL methods through an elementary convolution operator as described in [35]. As input, we provide the 3D coordinates of the atoms and the surface representation and atom types as a one-hot encoded vector. To study subtle differences between edge lengths across all proteins, we use Fourier distance features [36] defined by

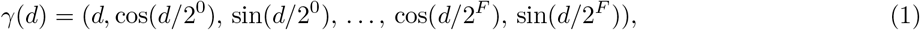

which are embedded together with the raw atomic information (i.e., nonlinearly transformed by an MLP). Although neural networks can learn complex features without feature engineering [37], we enhance this process by adding some essential features which are known to be relevant for the tasks: hydrophobicity and hydrogen bond potential. For this, we follow the preprocessing pipeline described in [28], where for every surface patch containing this raw atomic information, we determine and quantify these features.

#### 2.3.2 Geometric features

An important feature that becomes available when assuming a sufficiently smooth surface is curvature. We assume that the shape of surface patches, quantified by curvature features, plays an essential role in identifying interaction sites (see [38] and the references therein). In the current literature, two approaches of incorporating curvature are prevalent. The first approach relies on the Gaussian [39] and mean curvatures [40], intrinsic and extrinsic measures of curvature, respectively. The second approach employs the so-called shape index and the curvedness, which reflect the local shape of a surface patch and problem-specific length scales [41]. Both methods depend on the principal curvatures k_1_ and k_2_ [42], i.e., minimal and maximal normal curvatures, which are the eigenvalues of the shape operator [43]

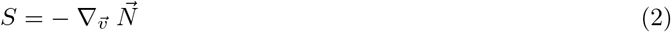

on the tangent basis. Intuitively, S measures the deviation of the normal vector field 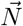 with respect to an arbitrary tangent vector 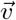. As is typical when dealing with meshes, the normal vertex vectors are estimated by averaging over the neighboring face normal vectors, computed using standard trigonometric procedures from the edges of the mesh.

To compute S, we employ the strategy introduced in [44]. We pick k nearest neighbors of a mesh node and project the relative vectors and the differences of the normal vectors into its tangent plane. First, a local basis needs to be constructed from the normals. Subsequently, we project the relative coordinates and relative normals to the tangent plane, and the shape operator S can be estimated.

In contrast to the two approaches described above, we do not rely on the principal curvatures’ explicit function(s). However, we do not use the raw geometric information imprinted in the principal curvatures since these quantities are coordinate-dependent. Thus, we diagonalize S and provide the eigenvalues, i.e., the principal curvatures k_1_ and k_2_, as input to a shallow MLP, which learns a latent representation suitable for downstream use.

### 2.4 Main DGRL pipeline

The main DGRL [29] pipeline processes the chemo-geometric embeddings using the 3D coordinates and surface normals as additional input (Figure 2E). It performs nonlinear geometry-informed processing of the features, resulting in a binary classification output. The underlying concept of this pipeline is weight sharing, i.e., convolutions. Most properties of an atom or surface point depend on its immediate neighborhood. Assuming that points in a neighborhood interact similarly across different neighborhoods, the choice of convolutional layers is natural to calculate the low dimensional representations of these properties. While our focus is the classification of binding sites and PPIs, this pipeline could also be used to learn other target values (e.g., a docking score).

The input of this processing step consists of the 3D coordinates and surfaces normals of the surface points under consideration, as well as the chemo-geometric features computed at those points. At each location, the features of all nearby points within a fixed radius are weighted and aggregated (2E). For a particular location, the weights describe the influence of all positions within the considered radius and are computed by an MLP, which maps the differences of point positions and surface normals to a scalar. As the MLP is applied to all possible point pairings, this operation step corresponds to a spatial convolution. The weights are then applied to the chemo-geometric features (transformed by an MLP), which are aggregated at each point by summation over the weighted features of all neighboring points. The diameter of the sphere results in 𝒪 (100-1000) neighbors per query point; thus, there are thousands of pairs per protein molecule, resulting in prohibitively large memory consumption if conventional methods are used. This problem is circumvented through the use of processing with symbolic operations, as provided by the PyKeOps library [45].

Finally, the aggregated features at each point are processed by a final MLP, resulting in an 8-dimensional feature vector per point, which is mapped to a binary output in the case of binding site prediction. For the interaction prediction task (Figure 2B), the outputs of the two proteins are further processed by another MLP, mapping to a binary output.

### 2.5 Model training

To label the surface meshes into a binding and non-binding vertices and aggregate evaluation metric of the predictions, we followed a similar strategy as described in [28]. For the interaction site prediction, all possible interaction sites were registered. For the PPIs prediction, we used an equal number of positive labels (i.e., surface patches on both proteins where they interact) and negative labels (i.e., surface patches on both proteins where they do not interact) to ensure a balanced dataset. The number of these discrete surfaces is determined by the size of the k-NN ball query.

We trained our models (i.e., the trainable parameters of the various MLPs, the ClusterGCN layer, etc.) using stochastic gradient descend with the AMSGrad version of the Adam optimizer [46]. We used the binary cross-entropy between the model output and the true target values as a loss function. To avoid overfitting, we employed an early stopping strategy [37], for which the model was evaluated on a validation set (10% of the training set) during training.

To evaluate our models, we followed [25] and used the AUROC (area under the receiver operating characteristic curve [47]) as evaluation metric. The final AUROC value summarizes predictions across all surface patches and quantifies the ratio of true positive rate vs. true negative rate for different cut-off values. For an example ternary complex (6W7O, Figure 3A), the AUROC is visualized in Figure 3B. As the final AUROC score for a protein-protein pair includes averaging over multiple positive and negative surface patch predictions involved in the PPIs, a high AUROC score for a single protein-protein interaction means that the model can accurately predict whether proteins will interact, based on the predictive power on individual surface patches of the two proteins.

**Figure 3:**
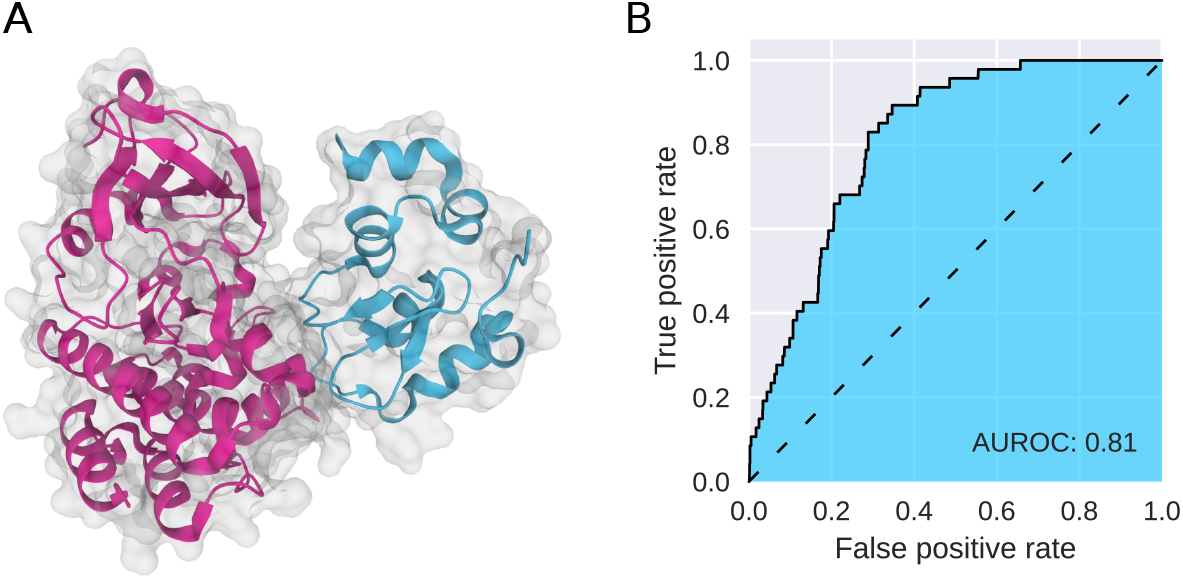
Example evaluation on ternary complex data. (A) PPIs of proteins involved in the ternary complex 6W7O (PDB ID). (B) Corresponding AUROC curve. See Appendix A for predictions on additional ternary complex data A.

### 2.6 Implementation

Our model was implemented using PyTorch [48], PyTorchGeometric [34], and PyKeOps [45].

## 3 Results

We present in this work a model capable of predicting possible interaction sites and whether two proteins will interact. As our goal is to perform these prediction tasks in the context of targeted protein degradation, we emphasize achieving good results on existing data of ternary complexes (consisting of two proteins and a degrader molecule). We first present the results of our model on two protein-protein interaction (PPI) datasets before discussing this particular scenario.

### 3.1 The *Orthogonal* dataset

To train and evaluate our model, we use two datasets. The first one, commonly called the MaSIF dataset, was used to evaluate the MaSIF model [28], an existing approach for PPI prediction, and consists of protein pairs from several different sources, including PDB [28]. As implemented in the MaSIF source code, in preprocessing, the MaSIF utilizes a distance cut-off value of 1.4 Å to assess if two surface points on different side chains are interacting or not. Since the model employs surface point clouds representation as inputs, we reasoned that having protein structures with higher crystallographic resolution could lead to more precise predictions, especially for identifying interaction sites. Furthermore, a more diverse dataset allows testing the generalizability of PPI prediction approaches, resulting in more robust results. We thus generated a new dataset by selecting all Homo sapiens proteins within PDB with a resolution of less than or equal to 2 Å, which were not included in the MaSIF dataset. The protein pairs in the resulting dataset, which we call *Orthogonal*, have an in average resolution of 1.7 Å (see Table 1).

**Table 1:**
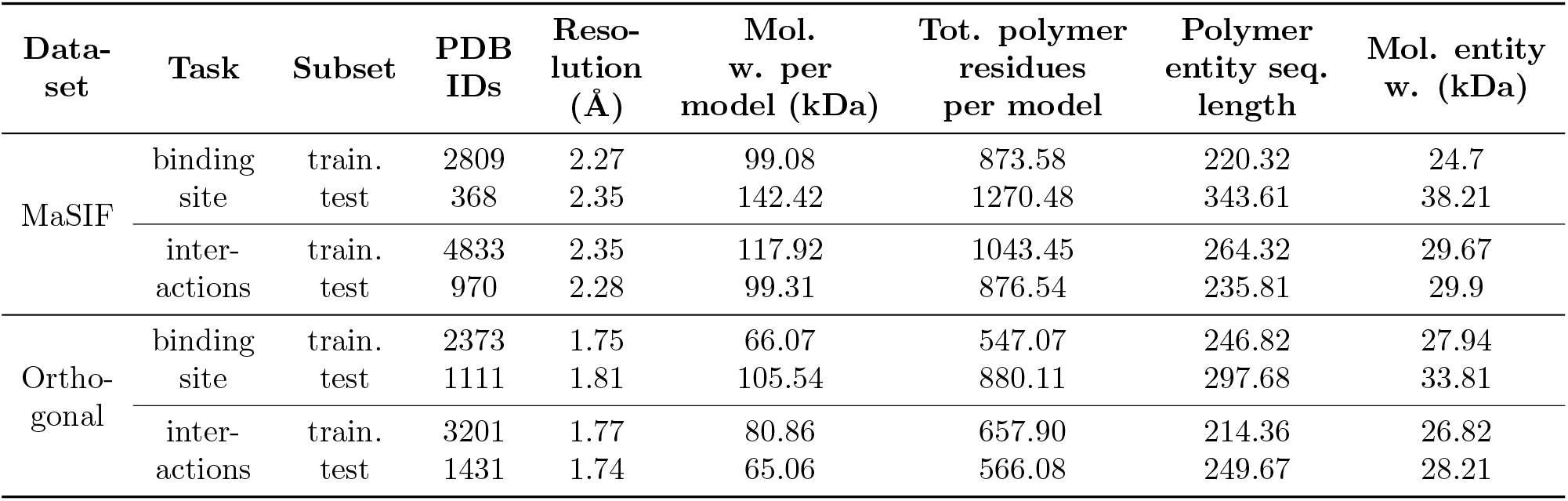
Comparison between the MaSIF and Orthogonal datasets. Columns show (from left to right) the dataset name, the considered task (prediction of binding sites or protein-protein interactions), the subset (training or test set), the number of PDB IDs within the datasets, the average resolution of the PDB ids within the datasets in Å, the average molecular weight per deposited model, the average total number of polymer residues per deposited model, the average polymer entity sequence length, and the average molecular entity weight.

We processed the selected protein’s FASTA sequences to the InterPro software [49], which categorized their structures into protein superfamilies and assigned a similarity score to each protein. The similarity scores were used to generate training and testing datasets that included a balanced representation of all protein classes and families. The main reason for the sequencing analysis is that we aim to obtain as much structural variability as possible to make the final model less susceptible to out-of-distribution protein structures.

It was necessary to visually examine all of the PDB structures in the orthogonal dataset to determine which chains of each PDB structure represent in a binary protein-protein interaction. We selected two chains for each PDB ID that reflects the PPI based on the proximity of chains 3D structures. Based solely on visual examination, we also speculated that the proteins in the orthogonal dataset had a higher number of tertiary protein structures than MaSIF, which could assist the model in being more generalizable and tolerating to surface variance of the proteins. This property is critical because tertiary protein structures are found in practically all highly complex biological systems, including transcription factors, signaling proteins, and membrane proteins, all of which are critical for drug discovery and development.

In total, the Orthogonal dataset contains 2373 and 1111 distinct PDB IDs for training and testing our model of binding site prediction, respectively, and 3201 and 1431 PDB IDs for training and testing of the protein-protein interaction predictions. As the ratio of test data to training data is very high, evaluation results on the test set of this dataset are likely to be a more accurate and reliable indicator of the generalization capabilities of a model compared to the dMaSIF dataset (where the ratio is lower). Furthermore, the dataset also includes rare complex protein residues that have heavy atoms bound to the side chain of the amino acids and operate as an organometallic bridge, providing more diverse data for further improvement of the evaluation accuracy.

### 3.2 PPI prediction on protein pairs

We trained and evaluated our model (see Section 2) using these two datasets and compared it to two prominent PPI prediction models (Table 2): MaSIF and dMaSIF (note that MaSIF refers to both a model and a dataset). For the MaSIF dataset, our model shows better results on the interaction prediction task, while dMaSIF results in higher accuracy on the binding site prediction task. While the former is arguably more critical for our goal of predicting PPIs for targeted protein degradation, we overall found no significant improvement on this dataset. Conversely, when evaluating the models on the Orthogonal dataset, we find that our model clearly outperforms dMaSIF (Table 2). For the task of interaction prediction in particular, there is a significant increase of the AUROC from 0.77 (dMaSIF) to 0.88 (our model). As the Orthogonal dataset includes more diversity, we conclude that our model can generalize better to a wide range of protein-protein pairs. Note that we used the default configuration (3 layers, patch size 12 Å) for the dMaSIF model, and it might be possible to improve its results on this dataset through hyperparameter tuning.

**Table 2:** Evaluation of PPI prediction models. The models’ AUROC scores were evaluated on the test set of the MaSIF dataset and our new Orthogonal dataset for the two tasks of binding site and interaction prediction. While our model does not significantly improve the results of dMaSIF [25] on the MaSIF dataset (with better results for interaction prediction but inferior results for binding site prediction), it significantly outperforms dMaSIF on the new Orthogonal dataset, which better captures the diversity of possible protein binding pairs.

| Dataset | Task | MaSIF<br>[28] | dMaSIF<br>[25] | Ours |
| --- | --- | --- | --- | --- |
| MaSIF | binding site | 0.85 | **0.87** | 0.82 |
|  | interactions | 0.81 | 0.82 | **0.88** |
| Orthogonal | binding site | - | 0.77 | **0.79** |
|  | interactions | - | 0.77 | **0.88** |

### 3.3 Evaluation on ternary complex data

We now turn to ternary complex data to evaluate the usefulness of our model for the development of new drugs for targeted protein degradation. In particular, we are interested in recovering the protein-protein interactions involved in ternary complexes using our model. A high recovery rate indicates that our model captures the essential features underlying ternary complex formation and thus can be used to evaluate previously unconsidered protein pairs to find new potential ternary complexes.

Following [51], we evaluated our trained model on 16 ternary complexes (listed in Table 3). The preprocessing of the proteins involved in the ternary complexes was identical to the general PPI prediction tasks (see Section 2), i.e., it included the construction of molecular surface meshes. We then processed the input data with our model pipeline, generating chemo-geometric features and DGRL embeddings, which were then used to make binary predictions. As this pipeline cannot account for the presence of the degrader molecule in the ternary complex, we ignored them in this analysis.

**Table 3:** Ternary complex dataset. In addition to the unique PDB identifiers and the chains, we also quote the name of E3 ligases and targets/POI (and domain of binding if applicable) [50, 51]. Note that for the PDB identifier 6W8I, three distinct pairs of chains are considered, which define BTK bound to the BIR3 domain of cIAP1.

| PDB ID | Chains | E3 ligase | Target |
| --- | --- | --- | --- |
| 5T35 | A, D | VHL | BRD4 BD2 |
| 6BN7 | B, C | CRBN | BRD4 BD1 |
| 6BN8 | B, C | CRBN | BRD4 BD1 |
| 6BN9 | B, C | CRBN | BRD4 BD1 |
| 6BNB | B, C | CRBN | BRD4 BD1 |
| 6BOY | B, C | CRBN | BRD4 BD1 |
| 6HAX | A, B | VHL | SMARCA2 |
| 6HAY | E, F | VHL | SMARCA2 |
| 6HR2 | E, F | VHL | SMARCA4 |
| 6SIS | E, H | VHL | BRD4 BD2 |
| 6W7O | B, D | cIAP1-BIR3 | BTK |
| 6W8I | A, D \| B, E \| C, F | cIAP1-BIR3 | BTK |
| 6ZHC | AD | VHL | Bcl-xL |
| 7KHH | CD | VHL | BRD4 BD1 |

We evaluated the interaction prediction AUROC on the ternary complex data for our model, trained on different datasets (Figure 4. When trained on the MaSIF dataset, the mean AUROC is 0.75, and the histogram of AUROCs is bimodal, with a few very low AUROC scores (Figure 4A). From this, we conclude that some of the proteins within the ternary complex dataset are very dissimilar to the data contained in the MaSIF dataset (i.e., they are out of distribution). Thus, the predictions on these proteins are very poor. We next evaluated the Orthogonal dataset (Figure 4B). Here, the mean AUROC has improved to 0.80, and there are no AUROC scores < 0.6. We thus conclude that the Orthogonal dataset with its larger diversity (see above) better reflects the proteins involved in the ternary complexes, and thus, this dataset is more useful for models involved in the prediction of ternary complexes.

**Figure 4:**
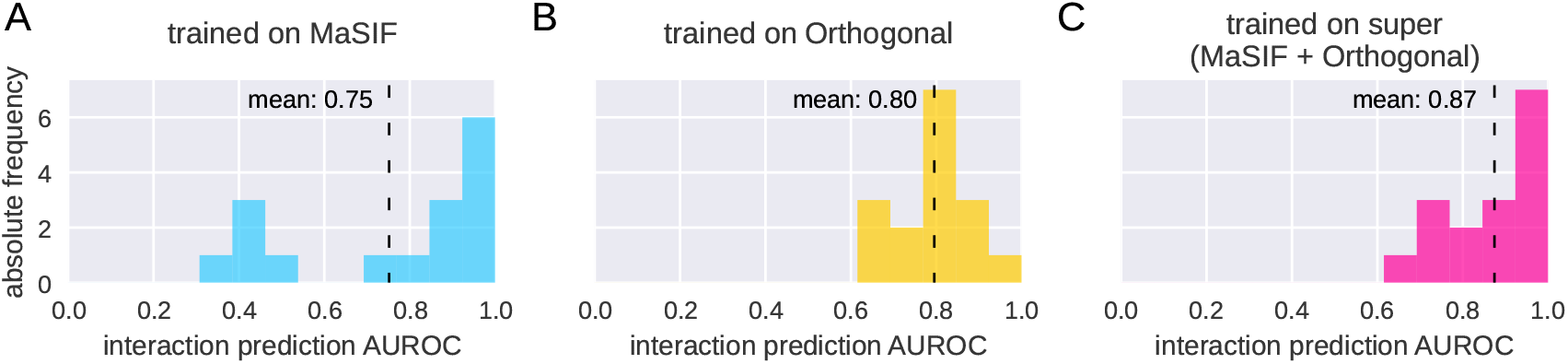
Histograms of AUROC of PPIs prediction for the ternary complex dataset defined in Table 3, for models trained on different training sets. (A) Results for a model trained on the MaSIF dataset. (B) Results for a model trained on the Orthogonal dataset. (C) Results for a model trained on the super dataset, i.e., the union of the MaSIF and Orthogonal datasets. The increase of the mean AUROC indicates that the MaSIF dataset lacks diversity of protein structures, thus, it is not sufficient for the development of models in the context of targeted protein degradation. Adding the structures of the Orthogonal dataset lead to a strong improvement. See Appendix A for the corresponding raw data.

When comparing Figures 4A and B, we find that though the mean has improved, the number of very high AUROC scores has decreased. We hypothesized that part of the MaSIF dataset is, in fact, very useful for the prediction of PPIs on the ternary complex data and that training jointly on both datasets could improve the predictions. We therefore re-trained our model on this “super”-dataset (MaSIF + Orthogonal), consisting of 8151 protein pairs for training and 2411 for evaluation. As expected, this lead to an even higher mean AUROC on the ternary complex data (0.87 instead of 0.80 and 0.75 individually, Figure 4). This provides further evidence that for the prediction of PPIs within a ternary complex, models require datasets that include the largest available diversity.

From the high accuracy of this final model, we conclude that the combination of the proposed geometric deep learning architecture paired with the novel dataset can accurately determine whether interactions between two proteins involved in the ternary complexes will take place. To demonstrate the ability of the algorithm to detect the optimal binding site, Figure 5 shows the surface point clouds that are hypothesized to determine the binding sites of both chains of the 6BN8 ternary complex (without degrader). We find that the algorithm can properly detect both the binding sites of each protein and the main interaction site between the two proteins.

**Figure 5:**
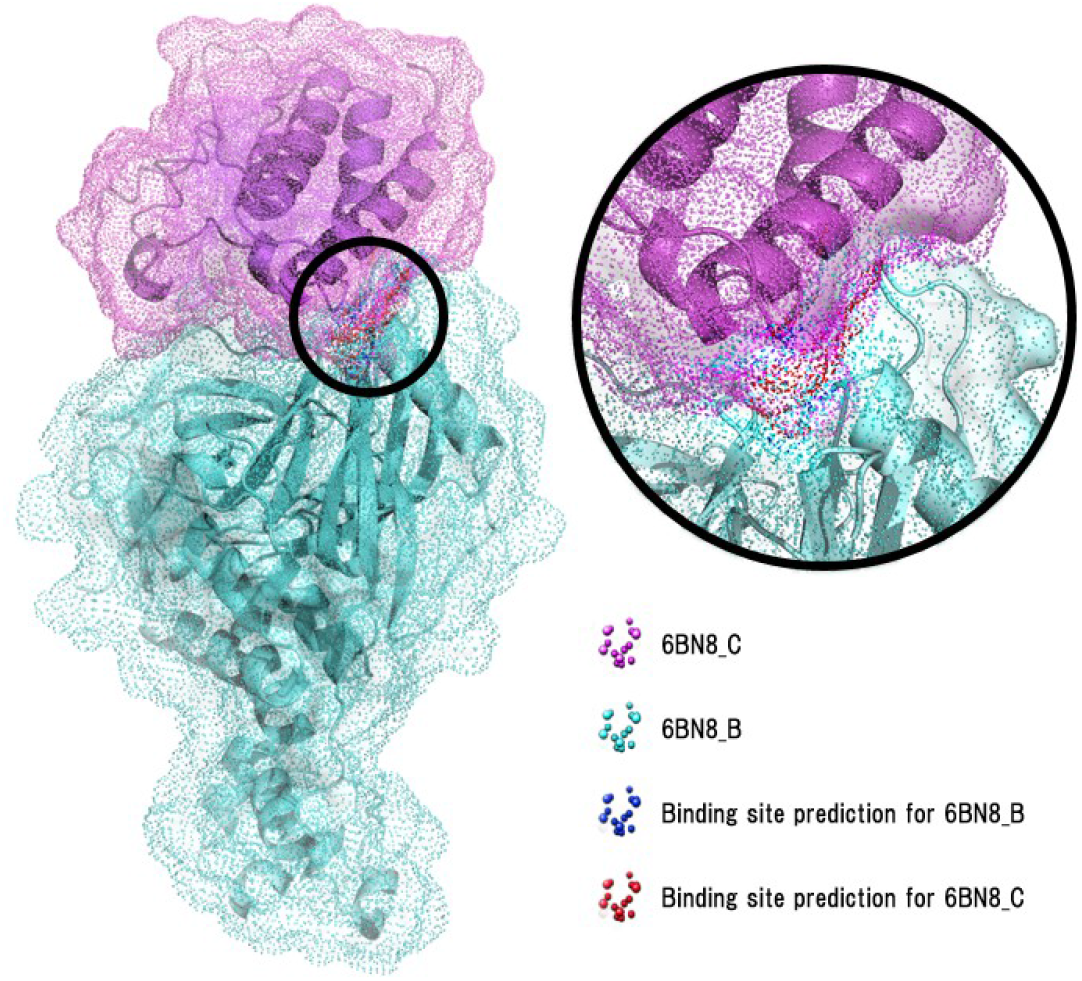
Biochemical representation of predictions for the ternary complex 6BN8 (PDB ID). Surface point clouds are superimposed to the original PDB structure. Blue and red points indicate interacting points prediction for chains B and C, respectively (AUROC: 0.987, inference performed using the model trained on the “super” dataset).

Overall, these results suggest that PPI prediction with DGRL is a promising tool for targeted protein degradation, and can be used, e.g., as a filtering mechanism to select possible protein-of-interest and E3 ligase pairs, which will, given a suitable degrader molecule, form a ternary complex.

## 4 Discussion

The machine learning model presented in this work used a novel DGRL pipeline based on surfaces meshes and was shown to be capable of predicting binding sites and interactions between proteins with high accuracy. We have furthermore presented a new, complex dataset that highlights the generalization capabilities of our model, demonstrating an improvement over previous work. Towards our primary goal of using protein-protein interaction (PPI) prediction for targeted protein degradation (TPD), we have demonstrated that our model can accurately predict the PPIs on the presently available ternary complex data.

### Related work

We showed that our model reaches results comparable to the MaSIF [28] and dMaSIF [25] models on the MaSIF dataset [28]. While dMaSIF, the current state-of-the-art model for PPI prediction based on 3D structural information, reaches a higher AUROC for binding site prediction, our model performed better than dMaSIF on the task of predicting interactions between pairs of proteins. For our goal of using PPI prediction for TPD, for which a model should determine pairings of a target protein and an E3 ligase, the latter task can be considered more important, and thus, our results constitute an improvement over the dMaSIF model. Furthermore, although our model design differs from dMaSIF in several important ways (e.g., we process explicitly computed surface meshes), inference in our model is still rapid (a few seconds per pass on a Tesla V100 machine) thus making it possible to screen many possible candidate pairs.

A variety of models for PPI prediction have been recently proposed, with some also using the MaSIF dataset as a benchmark. The model presented in [52] achieved AUROC scores of *≤*0.71 on this dataset, while is less than both dMaSIF and our model. The model presented in [53] was based on segmentation, which is problematic if only small portions of the protein surfaces overlap. Therefore, the authors used a modified version of the MaSIF dataset, rendering the results incomparable to ours [53]. While these models are based on the prediction of PPIs based on 3D structural information (template-based approaches), several models (e.g., [54, 55, 24]) follow the template-free paradigm and aim to predict PPIs only based on sequence information, with final results which are hard to compare to structure-based models as different datasets and evaluation metrics are used. In some cases, these models employ similar graph-based techniques as our model (e.g., [55]), highlighting their usefulness. We note that since the prediction of protein structure from sequence information has been approached experimental accuracy [56], we expect that these different approaches will converge in the future. This may result in models which use the best of both worlds by first predicting the 3D structure and then incorporating the various techniques (e.g., DGRL or 3D convolutions [57]) used to predict PPIs from structural information.

### The importance of diverse datasets for PPI prediction

It is well-known in machine learning that adequate datasets are required for models to be useful. In particular, the dataset used for training a model should cover all the relevant diversity of possible inputs. If this is not the case, then the model will not perform well on new, unseen data (i.e., it will not generalize well). This highlights the benefit of our new, more diverse Orthogonal dataset: as it includes protein pairs with significant differences to those within the MaSIF dataset, testing models on this dataset results in a more accurate estimate of the expected performance of a model on real-world data. Thus, the significantly higher performance of our model on this dataset suggests that it better captures the relevant features of diverse proteins than the dMaSIF model.

This is also corroborated by the analysis of our model on the ternary complex data, which revealed that adding the data from the Orthogonal dataset to the training data dramatically increases the prediction results. This shows that our new dataset is particularly valuable for PPI prediction in this context. We note that the dataset we employed here only contained positive examples of proteins involved in the ternary complex. For a better estimate of how well models can predict PPIs within ternary complexes, enhanced datasets with decoys (i.e., protein pairs with no interactions) as well as more data (as it becomes available) should be used in future works.

### Using PPI prediction for targeted protein degradation

As we have shown, training on a dataset that includes real-world diversity is crucial for machine learning models to be used in the context of targeted protein degradation. Our results suggest using a PPI prediction model as part of (e.g.) a filtering step in a preliminary screening process, where potentially interacting POIs and E3 ligases are identified for subsequent in-depth analysis. Such an approach can utilize the benefits of machine learning-driven methods to limit the number of potential candidates which need to be evaluated in computationally costly docking simulations or in time-and resource-intensive lab experiments.

One drawback of the current approach is that the presence of the degrader molecule is not taken into account. As there are three separate entities underlying a ternary complex, the PPIs there do not represent classical binary PPIs. Neglecting the degrader nevertheless constitutes a reasonable approximation when using PPI prediction as a proxy to predict potential ternary complexes, as the PPIs between the POI and the E3 ligase are an essential factor underlying ternary complex stability (Figure 1C; see, e.g., [18, 58]). For a ternary complex to be formed, positive cooperativity (i.e., PPIs) between the proteins is required, otherwise, only binary complexes (i.e., the degrader binding only to individual proteins) are formed. The approach presented here is based on the assumption that the factors underlying PPIs within ternary complexes are essential the same as in classical binary PPIs, e.g., portions of the protein surface which do not interact favorably with the solvent (water) are more likely to allow PPIs. However, future work should also explicitly take into account the presence of the degrader molecule, in particular, the physical constraints it imposes on the interaction of the two proteins. Recent machine learning approaches like PhysNet [59] or DimeNet [60] would be well-suited to endow models like dMaSIF or the one proposed in this work with the required spatial knowledge for generating (if possible) a degrader conformation. Another aspect that should be taken into account is the interactions between the degrader molecule and the proteins, for which the prediction of possible binding sites (as provided by our model or dMaSIF) may be useful.

Apart from targeted protein degradation with bifunctional degraders, there are also molecular glues [61], for which a monovalent small molecule alters the conformation of a protein in a way that enables PPIs. In this case, if the modified structures are available, PPI prediction (without taking into account the degrader) is enough to predict the efficacy of the degradation, and models such as the one presented in this work should allow predicting potential POI-E3 ligase pairs.

## Acknowledgments

We thank Michael Brunsteiner and Hosein Fooladi for helpful comments on this manuscript.

## Appendix

### A Prediction results for the known ternary complexes

**Table 4:**
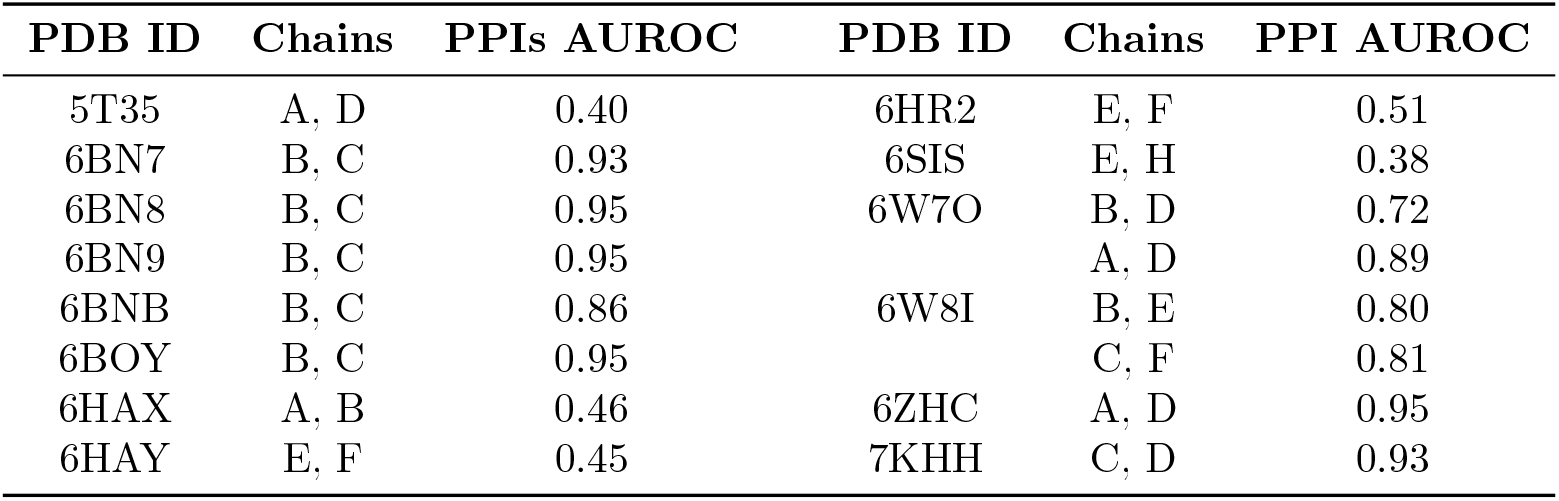
AUROC scores for PPIs prediction of our model trained on the MaSIF dataset shown in the Panel A of Figure 4.

**Table 5:**
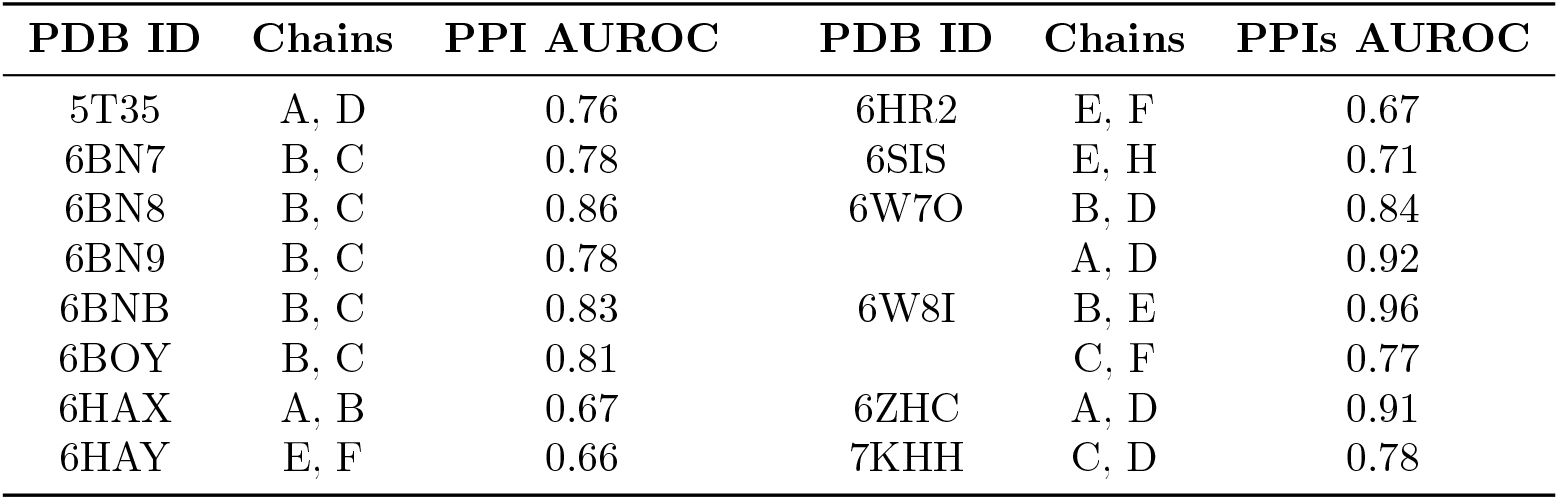
AUROC scores for PPIs prediction of our model trained on the orthogonal dataset shown in Panel B of Figure 4.

**Table 6:** AUROC scores for PPIs prediction of our model trained on the super-dataset shown in the Panel C of Figure 4.

| PDB ID | Chains | PPIs AUROC | PDB ID | Chains | PPI AUROC |
| --- | --- | --- | --- | --- | --- |
| 5T35 | A, D | 0.76 | 6HR2 | E, F | 0.69 |
| 6BN7 | B, C | 0.98 | 6SIS | E, H | 0.72 |
| 6BN8 | B, C | 0.99 | 6W7O | B, D | 0.81 |
| 6BN9 | B, C | 0.97 |  | A, D | 0.86 |
| 6BNB | B, C | 0.94 | 6W8I | B, E | 0.87 |
| 6BOY | B, C | 0.98 |  | C, F | 0.91 |
| 6HAX | A, B | 0.84 | 6ZHC | A, D | 0.97 |
| 6HAY | E, F | 0.74 | 7KHH | C, D | 0.97 |

## Notes

### Competing Interest Statement

The authors have declared no competing interest.

## References

[1] Daniel E. Koshland. The key-lock theory and the induced fit theory. Angewandte Chemie, 33:2375–2378, 1995.

[2] A. Hopkins and C. Groom. The druggable genome. Nat Rev Drug Discov, 1:727–730, 2002.

[3] John P. Overington, Bissan Al-Lazikani, and Andrew L. Hopkins. How many drug targets are there? Nat Rev Drug Discov, 5:993–996, 2006.

[4] Rita Santos, Oleg Ursu, Anna Gaulton, A. Patrícia Bento, Ramesh S. Donadi, Cristian G. Bologa, Anneli Karlsson, Bissan Al-Lazikani, Anne Hersey, Tudor I. Oprea,, and John P. Overington. A comprehensive map of molecular drug targets. Nat Rev Drug Discov, 1:19–34, 2017.

[5] Te-Wen Lo et al. Precise and heritable genome editing in evolutionarily diverse nematodes using TALENs and CRISPR/Cas9 to engineer insertions and deletions. International Journal of Proteomics, 195(2):331–48, 2013.

[6] Wei Xu, Xuezhen Jiang, and Linfeng Huang. RNA Interference Technology. Comprehensive Biotechnology, pages 560–575, 2019.

[7] Mingming Wei et al. First orally bioavailable prodrug of proteolysis targeting chimera (PROTAC) degrades cyclin-dependent kinases 2/4/6 in vivo. European journal of medicinal chemistry, 209, 2021.

[8] Stuart L. Schreiber Christopher J. Gerry. Unifying principles of bifunctional, proximity-inducing small molecules. Nat Chem Bio, 16:369–378, 2020.

[9] Sachini U Siriwardena, Dhanushka NP Munkanatta Godage, Veronika M Shoba, Sophia Lai, Mengchao Shi, Peng Wu, Santosh K Chaudhary, Stuart L Schreiber, and Amit Choudhary. Phosphorylation-inducing chimeric small molecules. Journal of the American Chemical Society, 142(33):14052–14057, 2020.

[10] Sayumi Yamazoe, Jeffrey Tom, Yue Fu, Wenqiong Wu, Liang Zeng, Changlei Sun, Qi Liu, Jie Lin, Kui Lin, Wayne J Fairbrother, et al. Heterobifunctional molecules induce dephosphorylation of kinases–a proof of concept study. Journal of medicinal chemistry, 63(6):2807–2813, 2019.

[11] Wesley W Wang, Li-Yun Chen, Jacob M Wozniak, Appaso M Jadhav, Hayden Anderson, Taylor E Malone, and Christopher G Parker. Targeted protein acetylation in cells using heterobifunctional molecules. Journal of the American Chemical Society, 143(40):16700–16708, 2021.

[12] Kathleen M. Sakamoto, Kyung B. Kim, Akiko Kumagai, Frank Mercurio, Craig M. Crews, and Raymond J. Deshaies. Protacs: Chimeric molecules that target proteins to the skp1–cullin–f box complex for ubiquitination and degradation. Proceedings of the National Academy of Sciences, 98(15):8554–8559, 2001.

[13] Aaron Ciechanover and Alan L. Schwartz. The ubiquitin-proteasome pathway: The complexity and myriad functions of proteins death. Proc. Natl. Acad. Sci. USA, 95, 1998.

[14] Ashton C Lai and Craig M Crews. Induced protein degradation: an emerging drug discovery paradigm. Nature reviews Drug discovery, 16(2):101–114, 2017.

[15] Dhanusha A. Nalawansha and Craig M. Crews. Protacs: An emerging therapeutic modality in precision medicine. Cell Chemical Biology, 27:998–1114, 2020.

[16] Mariell Pettersson and Craig M. Crews. Proteolysis targeting chimeras (protacs) — past, present and future. Drug Discovery Today: Technologies, 31:15–27, 2019. Protein degradation drug discovery.

[17] Miklós Békés, David R Langley, and Craig M Crews. Protac targeted protein degraders: the past is prologue. Nature Reviews Drug Discovery, pages 1–20, 2022.

[18] Scott J Hughes and Alessio Ciulli. Molecular recognition of ternary complexes: a new dimension in the structure-guided design of chemical degraders. Essays in biochemistry, 61(5):505–516, 2017.

[19] Tasuku Ishida and Alessio Ciulli. E3 ligase ligands for protacs: How they were found and how to discover new ones. SLAS DISCOVERY: Advancing the Science of Drug Discovery, 26(4):484–502, 2021. PMID: 33143537.

[20] Brandon Charles Seychell and Tobias Beck. Molecular basis for protein–protein interactions. Beilstein Journal of Organic Chemistry, 17(1):1–10, 2021.

[21] Koh Takeuchi, Kumaran Baskaran, and Haribabu Arthanari. Structure determination using solution nmr: Is it worth the effort? Journal of Magnetic Resonance, 306:195–201, 2019.

[22] Jean-Paul et al. Renaud. Cryo-em in drug discovery: achievements, limitations and prospects. Nature reviews. Drug discovery, 17(7):471–492, 2018.

[23] Sharon Sunny and PB Jayaraj. Protein–protein docking: Past, present, and future. The protein journal, pages 1–26, 2021.

[24] Richard Evans, Michael O’Neill, Alexander Pritzel, Natasha Antropova, Andrew W Senior, Timothy Green, Augustin žídek, Russell Bates, Sam Blackwell, Jason Yim, et al. Protein complex prediction with alphafold-multimer. Biorxiv, 2021.

[25] Freyr Sverrisson, Jean Feydy, Bruno E. Correia, and Michael M. Bronstein. Fast end-to-end learning on protein surfaces. In Proceedings of the IEEE/CVF Conference on Computer Vision and Pattern Recognition (CVPR), pages 15272–15281, June 2021.

[26] Andras Szilagyi and Yang Zhang. Template-based structure modeling of protein–protein interactions. Current Opinion in Structural Biology, 24:10–23, 2014. Folding and binding / Nucleic acids and their protein complexes.

[27] Helen M. Berman, John Westbrook, Zukang Feng, Gary Gilliland, T. N. Bhat, Helge Weissig, Ilya N. Shindyalov, and Philip E. Bourne. The Protein Data Bank. Nucleic Acids Research, 28(1):235–242, 01 2000.

[28] Pablo Gainza, Freyr Sverrisson, Federico Monti, Emanuele Rodolá, Michael M. Bronstein, and Bruno E. Correia. Deciphering interaction fingerprints from protein molecular surfaces using geometric deep learning. Nature Methods, 17:184–192, 2020.

[29] William L Hamilton. Graph representation learning. Synthesis Lectures on Artifical Intelligence and Machine Learning, 14(3):1–159, 2020.

[30] Sangsoo Lim, Yijingxiu Lu, Chang Yun Cho, Inyoung Sung, Jungwoo Kim, Youngkuk Kim, Sungjoon Park, and Sun Kim. A review on compound-protein interaction prediction methods: Data, format, representation and model. Computational and Structural Biotechnology Journal, 19:1541–1556, 2021.

[31] Lisa Torrey and Jude Shavlik. Transfer learning. In Handbook of research on machine learning applications and trends: algorithms, methods, and techniques, pages 242–264. IGI global, 2010.

[32] Dong Xu and Yang Zhang. Generating Triangulated Macromolecular Surfaces by Euclidean Distance Transform. PLoS ONE, 4(12):e8140, 2009.

[33] Dong Xu, Hua Li, and Yang Zhang. Protein Depth Calculation and the Use for Improving Accuracy of Protein Fold Recognition. Journal of Computational Biology, 20(10):805–816, 2013.

[34] Matthias Fey and Jan Eric Lenssen. Fast Graph Representation Learning with PyTorch Geometric. Proceedings of the International Conference on Learning Representations, 2019.

[35] Wei-Lin Chiang, Xuanqing Liu, Si Si, Yang Li, Samy Bengio, and Cho-Jui Hsieh. Cluster-GCN: An Efficient Algorithm for Training Deep and Large Graph Convolutional Networks. Proceedings of the 25th ACM SIGKDD International Conference on Knowledge Discovery and Data Mining, 2019.

[36] Hannes Stärk et al. 3DInfoMax improves GNNs for molecular property prediction. 2110.04126, 2021.

[37] Ian Goodfellow, Yoshua Bengio, and Aaron Courville. Deep Learning. MIT Press, 2016. http://www.deeplearningbook.org.

[38] Shuangye Yin, Elizabeth A. Proctor, Alexey A. Lugovskoy, and Nikolay V. Dokholyana. Fast screening of protein surfaces using geometric invariant fingerprints. Proc. Natl. Acad. Sci. USA, 109(39):16622–16626, 9 2009.

[39] Eric W. Weisstein. Gaussian Curvature (Wolfram MathWorld). https://mathworld.wolfram.com/GaussianCurvature.html. Accessed: 14-10-2021.

[40] Eric W. Weisstein. Mean Curvature (Wolfram MathWorld). https://mathworld.wolfram.com/MeanCurvature.html. Accessed: 14-10-2021.

[41] Jan J. Koenderink and Andrea J. van Doorn. Surface shape and curvature scales. Image and Vision Computing, 10(8):557–564, 1992.

[42] Eric W. Weisstein. Principal Curvatures (Wolfram MathWorld). https://mathworld.wolfram.com/PrincipalCurvatures.html. Accessed: 03-01-2022.

[43] Eric W. Weisstein. Shape Operator (Wolfram MathWorld). https://mathworld.wolfram.com/ShapeOperator.html. Accessed: 14-10-2021.

[44] Yueqi Cao, Didong Li, Huafei Sun, Amir H. Assadi, and Shiqiang Zhang. Efficient weingarten map and curvature estimation on manifolds. Machine Learning, 110:1319–1344, 2021.

[45] Benjamin Charlier et al. Kernel Operations on the GPU, with Autodiff, without Memory Overflows. Journal of Machine Learning Research, 22(74):1–6, 2021.

[46] Sashank J. Reddi, Satyen Kale, and Sanjiv Kumar. On the Convergence of Adam and Beyond. Proceedings of the International Conference on Learning Representations, 2019.

[47] Andrew P. Bradley. The use of the area under the roc curve in the evaluation of machine learning algorithms. Pattern Recognition, 30(7):1145–1159, 1997.

[48] Adam Paszke, Sam Gross, Soumith Chintala, Gregory Chanan, Edward Yang, Zachary DeVito, Zeming Lin, Alban Desmaison, Luca Antiga, and Adam Lerer. Automatic differentiation in PyTorch. NIPS-W, 2017.

[49] Matthias Blum et al. The InterPro protein families and domains database: 20 years on. Nucleic Acids Research, 2020.

[50] Daniel Zaidman, Jaime Prilusky, and Nir London. PRosettaC: Rosetta Based Modeling of PROTAC Mediated Ternary Complexes. J. Chem. Inf. Model., 60:4894–4903, 2020.

[51] Gaoqi Weng, Dan Li, Yu Kang, and Tingjun Hou. Integrative Modeling of PROTAC-Mediated Ternary Complexes. Journal of Medicinal Chemistry, 64(21):16271–16281, 2021.

[52] He Huang, Chengshi Zeng, and Xinqi Gong. Inter-protein contact map generated only from intra-monomer by image inpainting. In 2021 IEEE International Conference on Bioinformatics and Biomedicine (BIBM), pages 131–136. IEEE, 2021.

[53] Bowen Dai and Chris Bailey-Kellogg. Protein interaction interface region prediction by geometric deep learning. Bioinformatics, 37(17):2580–2588, 2021.

[54] Minli Tang, Longxin Wu, Xinyu Yu, Zhaoqi Chu, Shuting Jin, and Juan Liu. Prediction of protein–protein interaction sites based on stratified attentional mechanisms. Frontiers in Genetics, 12, 2021.

[55] Qianmu Yuan, Jianwen Chen, Huiying Zhao, Yaoqi Zhou, and Yuedong Yang. Structure-aware protein–protein interaction site prediction using deep graph convolutional network. Bioinformatics, 38(1):125–132, 09 2021.

[56] John Jumper, Richard Evans, Alexander Pritzel, Tim Green, Michael Figurnov, Olaf Ronneberger, Kathryn Tunyasuvunakool, Russ Bates, Augustin žídek, Anna Potapenko, et al. Highly accurate protein structure prediction with alphafold. Nature, 596(7873):583–589, 2021.

[57] Nicolas Renaud, Cunliang Geng, Sonja Georgievska, Francesco Ambrosetti, Lars Ridder, Dario F Marzella, Manon F Réau, Alexandre MJJ Bonvin, and Li C Xue. Deeprank: a deep learning framework for data mining 3d protein-protein interfaces. Nature communications, 12(1):1–8, 2021.

[58] David Zollman and Alessio Ciulli. Structural and biophysical principles of degrader ternary complexes. In Protein Degradation with New Chemical Modalities, pages 14–54. Royal Society of Chemistry, 2020.

[59] Oliver T Unke and Markus Meuwly. Physnet: A neural network for predicting energies, forces, dipole moments, and partial charges. Journal of chemical theory and computation, 15(6):3678–3693, 2019.

[60] Johannes Klicpera, Janek Groß, and Stephan Günnemann. Directional Message Passing for Molecular Graphs. Proceedings of the International Conference on Learning Representations, 2020.

[61] Shanique Alabi. Novel mechanisms of molecular glue-induced protein degradation. Biochemistry, 60(31):2371–2373, 2021.

